# Inference for epidemic models with time varying infection rates: tracking the dynamics of oak processionary moth in the UK

**DOI:** 10.1101/2021.12.09.471950

**Authors:** L E Wadkin, J Branson, A Hoppit, N G Parker, A Golightly, A W Baggaley

## Abstract

1. Invasive pests pose a great threat to forest, woodland and urban tree ecosystems. The oak processionary moth (OPM) is a destructive pest of oak trees, first reported in the UK in 2006. Despite great efforts to contain the outbreak within the original infested area of South-East England, OPM continues to spread.
2. Here we analyse data of the numbers of OPM nests removed each year from two parks in London between 2013 and 2020. Using a state-of-the-art Bayesian inference scheme we estimate the parameters for a stochastic compartmental SIR (susceptible, infested, removed) model with a time varying infestation rate to describe the spread of OPM.
3. We find that the infestation rate and subsequent basic reproduction number have remained constant since 2013 (with *R*_0_ between one and two). This shows further controls must be taken to reduce *R*_0_ below one and stop the advance of OPM into other areas of England.
4. *Synthesis.* Our findings demonstrate the applicability of the SIR model to describing OPM spread and show that further controls are needed to reduce the infestation rate. The proposed statistical methodology is a powerful tool to explore the nature of a time varying infestation rate, applicable to other partially observed time series epidemic data.

## 1 Introduction

Invasive pests, such as non-native insects, pose a threat to forest, woodland and urban tree ecosystems by damaging and killing trees and reducing biodiversity [1–3]. This threat has increased in recent years due to growth in international travel and trade [4] coupled with a changing climate driving the migration of species into new ecosystems [5]. The loss of biodiversity has a profound economic impact, through short to long term control measures and the impact on ecosystem services [6–8].

The oak processionary moth (OPM), *Thaumetopoea processionea*, is an invasive and destructive pest of oak trees, causing defoliation and making trees vulnerable to other stressors and pathogens. The larvae of OPM have poisonous hairs, containing a urticating toxin (thaumetopoein) which is harmful to human and animal health [9–12].

OPM was introduced to the UK through accidental imports on live oak plants, first reported in 2006. Up to 2010, the governmental policy was one of eradication [13, 14]. However, in 2011 it was decided that OPM was fully established in the South-East England area and so the government moved to a containment strategy, aiming to contain the OPM infestations within this original outbreak area [14]. In 2018, legislation was introduced to curb continuing imports through the Plant Health Order [15]. Despite the containment strategies, the extent of OPM has continued to spread with recent analysis suggesting an expansion rate of 1.7 km/year for 2006–2014, with an increase to 6 km/year from 2015 onwards [16]. The regions surrounding the current infection area are particularly climatically suitable [17] and so being able to predict and control the future dynamics of the OPM population is crucial to protect these areas.

Mathematical models provide a powerful tool for describing and predicting the spread of tree disease and pest infections [18–20]. For OPM, previous work has included using models from electric network theory to predict high risk regions [21] along with species distribution models to examine the spatial distributions of OPM [22] and the effects of climate change on its expansion [17]. Bayesian inference can be used to inform and evaluate these ecological mathematical models [23]. Previously, Bayesian approaches have been used to estimate key parameters in the spatio-temporal invasion of alien species [24], however, the techniques have yet to be applied to data for the spread of OPM.

Nevertheless, the Bayesian paradigm provides a natural mechanism for quantifying and propagating uncertainty in the model parameters and dynamic components. Consequently, Bayesian inference techniques have been ubiquitously applied in the broad area of epidemiology [see e.g. 25–27, for an overview].

In this paper we use data tracking the numbers and locations of OPM nests removed from oak trees as part of a control program in two parks in south London. We illustrate the use of statistical inference techniques for estimating the parameters for a classic SIR compartmental model [28, 29] consisting of susceptible, infested and removed states. To allow for intrinsic stochasticity in the spread of OPM, we use an ItÔ stochastic differential equation [30] representation of the SIR model. This is further modified via the introduction of a time varying infestation rate, as is necessary to capture the effect of unknown influences such as preventative measures [31]. Bayesian inference for the resulting model is complicated by the intractability of the observed data likelihood, and subsequently, the joint posterior distribution of the key quantities of interest (model parameters and dynamic components). We overcome these difficulties via a linear Gaussian approximation of the stochastic SIR model, coupled with a Markov chain Monte Carlo scheme [32] for generating samples from the joint posterior. These methods are outlined in Section 2 and detailed in the Supplementary Information, Sections S1 and S2, for use as a toolbox to apply to other ecological datasets. We use the parameters from the compartmental model to estimate a yearly *R*_0_ measure for OPM, analogous to the basic reproduction number for a pathogen [33], and estimate the OPM population in 2021.

## 2 Methods

In this section we present the observational time series data with a summary of the data collection methods (Section 2.1), the details of the stochastic SIR model (Section 2.2), and an outline of our statistical inference methods (Section 2.3). Further statistical details including the relevant algorithms are set out in the Supplementary Information (Sections S1 and S2).

### 2.1 Data

The data used is part of the national Oak Processionary Moth Survey. The OPM survey is conducted by the Forestry Commission as part of the governmental OPM control programme [34]. The University of Southampton (GeoData) provide analysis, support and hold the data on behalf of the Forestry Commission. We note that for the wider OPM control programme, the data is collected for operational needs and therefore there are limitations for research purposes. For example, the surveillance strategy between 2013 and 2020 focussed on monitoring the expansion of the outer edge of the known infested area. Thus, although the data provides a sufficient estimate of the outer expansion, the presence of nests in the central infection zone is likely underestimated. However, in this paper we use a sub-set of this data from surveys in Richmond and Bushy Parks in which the whole park area was surveyed each year, and the number of nests recorded accurately.

The data used in this study was obtained through the recording of OPM presence in Richmond and Bushy Parks in South-West London. For each of the years 2013–2020 it contains i) the eastings and northings of infested tree and ii) the number of OPM nests removed from each tree. The dataset consists of 8470 unique infested trees, with 1767 in Bushy Park and 6703 in Richmond Park. The locations of infested trees are shown across the two parks in Figure 1.

**Figure 1:**
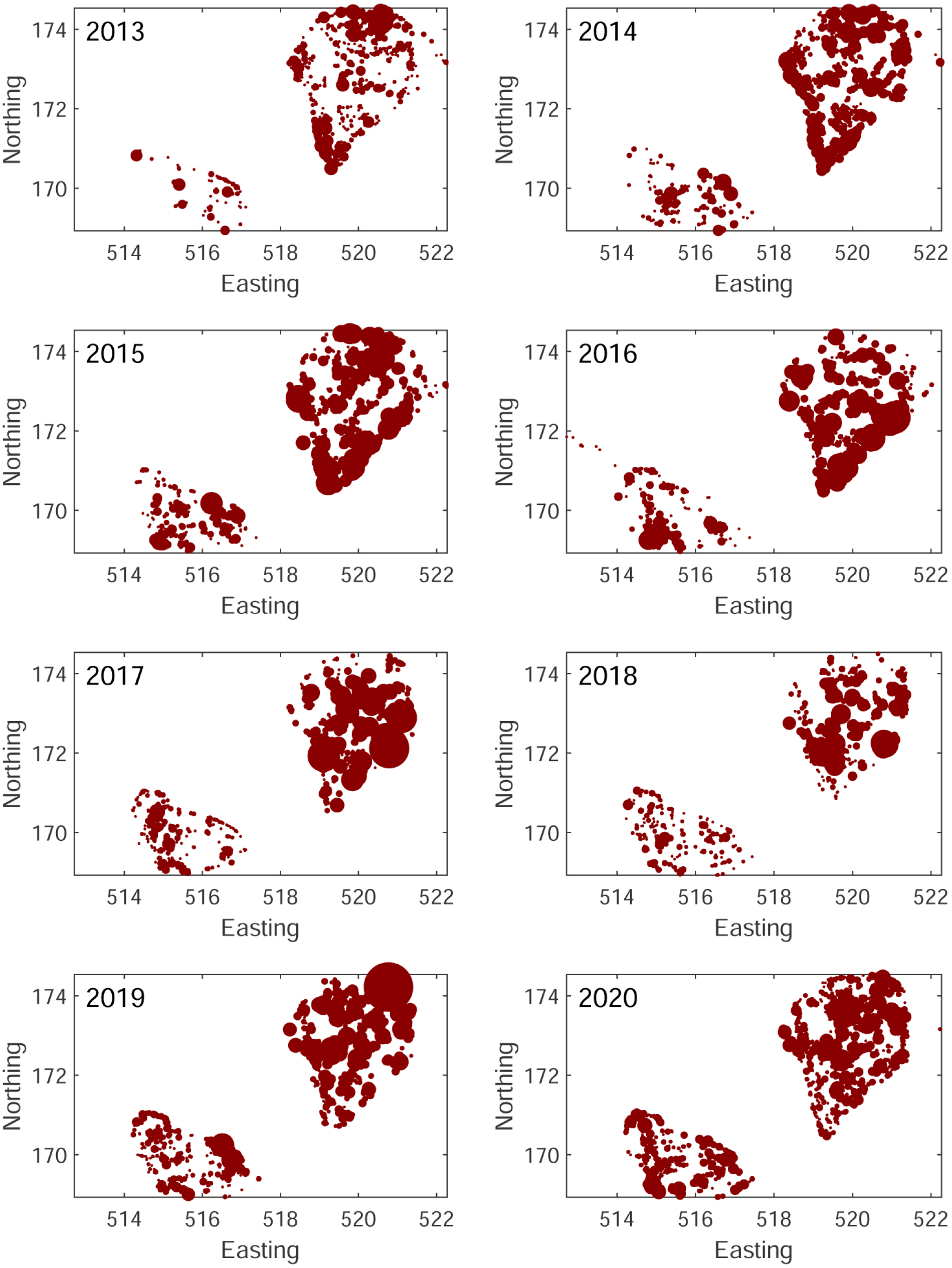
Map of the nests removed from Bushy (bottom left) and Richmond (top right) parks between 2013–2020. The area of the marker is proportional to the number of nests removed.

The raw and cumulative time series of the numbers of removed nests are shown in Figure 2(a) and (b). We count each infested tree in the year it was first infested as one ‘removal’ in the SIR model (see Section 2.2), regardless of how many nests were recorded as removed from this location. The raw and cumulative time series for the number of these removals is shown in Figure 2(c) and (d). We use the latter cumulative time series, *R*(*t*), as our observational data in the following sections.

**Figure 2:**
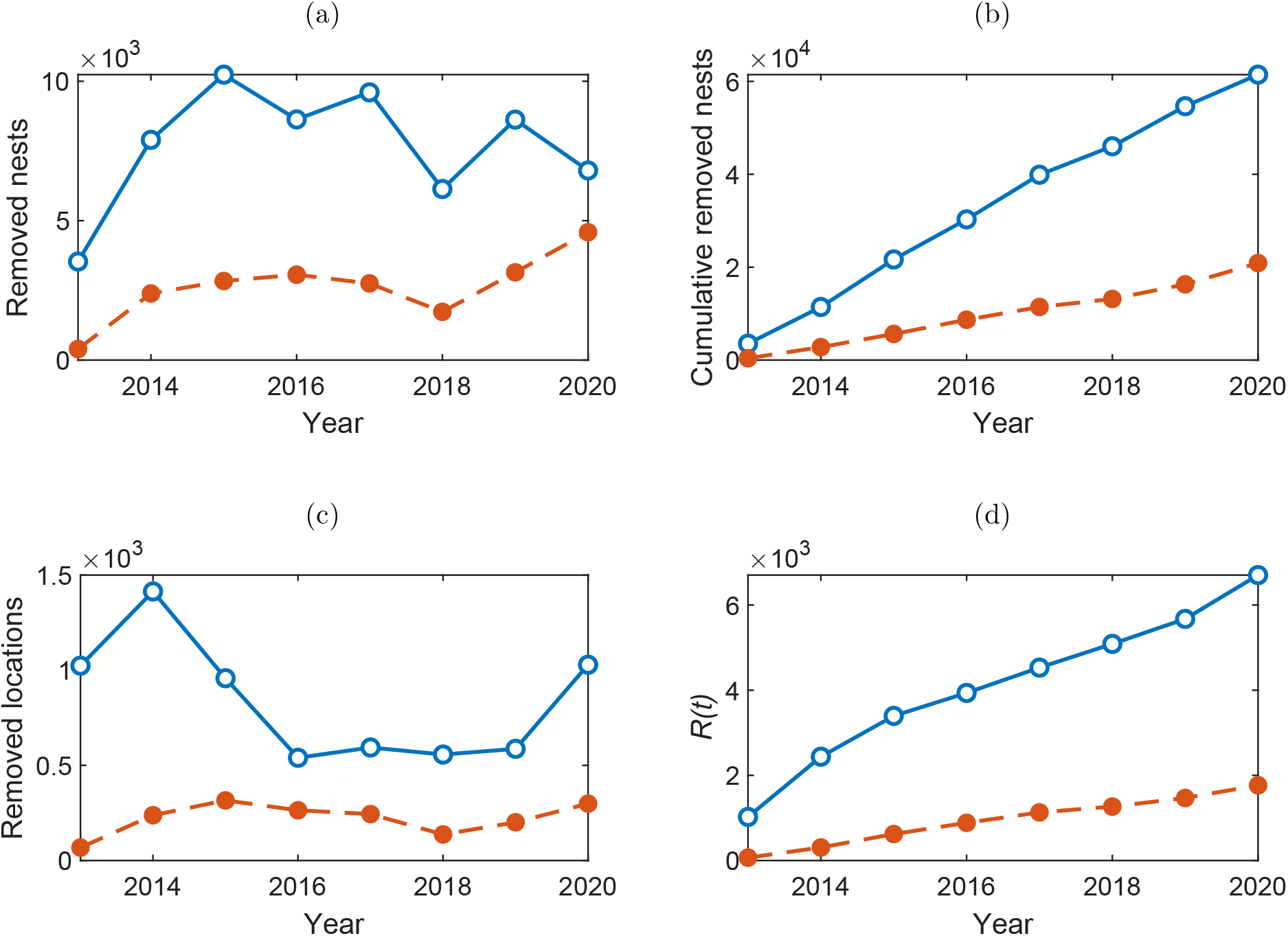
The (a) raw and (b) cumulative number of OPM nests removed from Richmond (blue) and Bushy (orange dashed) parks between 2013–2021. The number of (a) raw and (b) cumulative unique trees (described by their eastings and northings) which had nests removed between 2013–2021. The cumulative number of trees is the time series *R*(*t*) corresponding to the ‘removed’ category in the SIR model (see Section 2.2).

### 2.2 Stochastic SIR model

We consider an SIR model [28, 35] in which a population of fixed size *N* is classified into compartments consisting of susceptible (*S*), infected (*I*) and removed (*R*) individuals. Here we refer to the *I* compartment as ‘infested’ due to the context of describing an invasive pest presence. Transitions between compartments can be summarised via two pseudo-reactions of the form

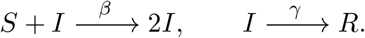

Hence, the first transition describes contact of an infective individual with a susceptible and with the net effect resulting in an additional infested individual and one fewer susceptible. The second transition accounts for removal (recovered with immunity, quarantined or dead) of an infested individual. The parameters *β* and *γ* govern the rate of infestation and removal, respectively. A fixed population of trees is appropriate as over the timescale of interest the number of trees born into the *S* compartment will be sufficiently small to be negligible. Our setting has individuals as trees and the contact process is understood to take place via the dispersal of OPM. It is clear that transitions should result in discrete changes to the numbers of trees in each state. This most naturally leads to a continuous time, discrete valued Markov jump process (MJP) description of disease dynamics, as detailed in the Supplementary Information, Section S1. We eschew the MJP formalism in favour of a continuous valued approximation, formulated as a stochastic differential equation (SDE). This is a pragmatic choice, since the SDE model ultimately leads to a computationally efficient inference scheme, and the model can be easily augmented with additional components, such as time varying parameters, which we now describe.

The SDE representation of the standard SIR model can be derived directly from the MJP (see Supplementary Information, Section S1). Here we extend this to include a time varying infection process. Let 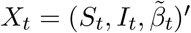 where *S*_*t*_ and *I*_*t*_ denote the numbers in each of the states *S* and *I* at time *t* ≥ 0 and 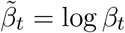 is the (transformed) time varying infection process. Note that the fixed population size gives *R*_*t*_ = *N* − *S*_*t*_ − *I*_*t*_ for all *t* ≥ 0 so that the current state of the SIR model is completely described by *X*_*t*_. We model 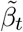 as a generalised Brownian motion process so that

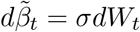

and *W*_*t*_ is a standard Brownian motion process. Hence we assume that the log infection rate evolves according to a random walk in continuous time, with variability controlled by *σ*. Combining this process with component SDEs describing the dynamics of *S*_*t*_ and *I*_*t*_ gives the complete SDE description of the SIR model with time varying infection rate as

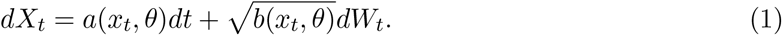

Here, *x*_*t*_ = (*s*_*t*_, *i*_*t*_, *β*_*t*_)′ is the state of the system at time *t*, *θ* = (*γ, σ*)′ is the vector of static parameter values, *W*_*t*_ = (*W*_1,*t*_, *W*_2,*t*_, *W*_3,*t*_)′ is a vector of uncorrelated standard Brownian motion processes, and the drift function *a*(*x*_*t*_, *θ*) and diffusion coefficient *b*(*x*_*t*_, *θ*) are given by

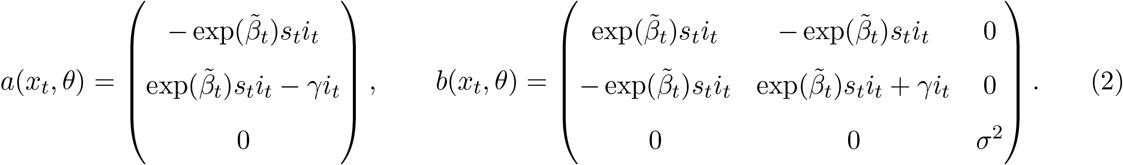

Unfortunately, due to the nonlinear forms of *a*(*x*_*t*_, *θ*) and *b*(*x*_*t*_, *θ*), the SDE specified by (1)–(2) cannot be solved analytically. We therefore replace the intractable analytic solution with a tractable Gaussian process approximation, which is the subject of the next section. The resulting linear noise approximation is subsequently used as the inferential model.

#### Linear noise approximation

The linear noise approximation (LNA) provides a tractable approximation to the SDE given by (1)–(2). In what follows we give a brief derivation; formal details can be found in [36] [see also 37, 38].

Consider a partition of *X*_*t*_ as

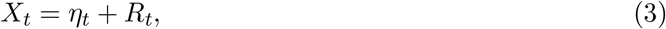

where {*η*_*t*_, *t* ≥ 0} is a deterministic process satisfying the ODE

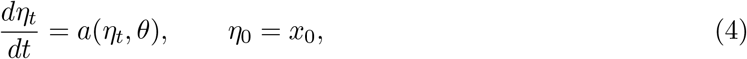

and {*R*_*t*_, *t* ≥ 0} is a residual stochastic process. The residual process *R*_*t*_ satisfies

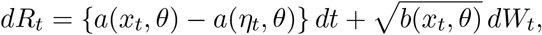

which will typically be intractable. Assumption that ||*X*_*t*_ − *η*_*t*_|| is “small” motivates a Taylor series expansion of *a*(*x*_*t*_, *θ*) and *b*(*x*_*t*_, *θ*) about *η*_*t*_, with retention of the first two terms in the expansion of *a* and the first term in the expansion of *b*. This gives an approximate residual process 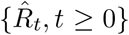 satisfying

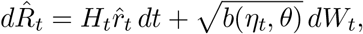

where *H*_*t*_ is the Jacobian matrix with (*i,j*)th element

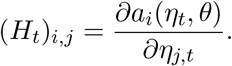

For the SIR model in (1)–(2) we therefore have

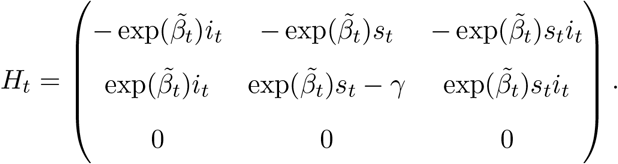

Given an initial condition 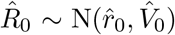, it can be shown that 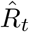 is a Gaussian random variable (see [32]). Consequently, the partition in (3) with *R*_*t*_ replaced by 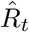, and the initial conditions *η*_0_ = *x*_0_ and 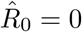 give

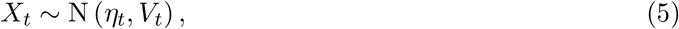

where *η*_*t*_ satisfies (4) and *V*_*t*_ satisfies

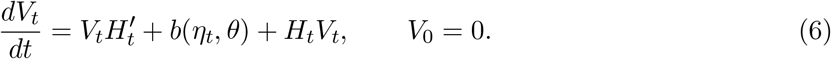

Hence, the linear noise approximation is characterised by the Gaussian distribution in (5), with mean and variance found by solving the ODE system given by (4) and (6). Although this ODE system will typically be intractable, a numerical scheme can be straightforwardly applied.

### 2.3 Bayesian inference

We consider the case in which not all components of the stochastic epidemic model are observed. Moreover, we assume that data points are subject to measurement error, which accounts for mismatch between the latent and observed process, due to, for example, the way in which the data are collected. Observations (on a regular grid) *y*_*t*_, *t* = 0, 1, . . . *n* are assumed conditionally independent (given the latent process) with conditional probability distribution obtained via the observation equation,

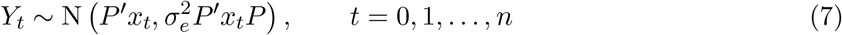

where *P* = (1, 1, 0)′. This choice of *P* is due to the data consisting of observations on the removed state *R*_*t*_, which, for a known population size *N*, is equivalent (in information content) to observing the sum *S*_*t*_ + *I*_*t*_. Note that the logarithm of the infection rate process, 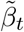 is completely unobserved. Our choice of observation model is motivated by a Gaussian approximation to a Poisson *Po*(*P*′*x*_*t*_*P*′) distribution, with the role of 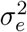 to allow a decoupling of the mean and variance. Moreover, the assumption of a Gaussian observation model admits a tractable observed data likelihood function, when combined with the LNA (see Section 2.2) as a model for the latent epidemic process *X*_*t*_. Details on a method for the efficient evaluation of this likelihood function can be found in Section S2.3 of the Supplementary Information.

Given data *y* = (*y*_0_, *y*_1_, . . ., *y*_*n*_) and upon ascribing a prior density *π*(*θ*) to the components of *θ* = (*γ, σ, σ*_*e*_)′ (augmented to include *σ*_*e*_), Bayesian inference proceeds via the joint posterior for the static parameters *θ* and unobserved dynamic process *x* = (*x*_0_, *x*_1_, . . ., *x*_*n*_). We have that

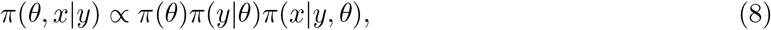

where *π*(*y*|*θ*) is the observed data likelihood and *π*(*x*|*y, θ*) is the conditional posterior density of the latent dynamic process. Although *π*(*y*|*θ*) and *π*(*x*|*y, θ*) can be obtained in closed form under the LNA, the joint posterior in (8) is intractable. In the Supplementary Information Section S2 we describe a Markov chain Monte Carlo scheme for generating (dependent) samples from (8). Briefly, this comprises two steps: i) the generation of samples *θ*^(1)^, . . ., *θ*^(*M*)^ from the marginal parameter posterior *π*(*θ*|*y*) ∝ *π*(*θ*)*π*(*y*|*θ*) and ii) the generation of samples *x*^(1)^, . . ., *x*^(*M*)^ by drawing from the conditional posterior *π*(*x*|*y, θ*^(*i*)^), *i* = 1, . . ., *M*.

Given inferences on the static parameters *θ* and the latent dynamic process *x*, we consider the following diagnostics for assessing model fit. The within sample predictive density is

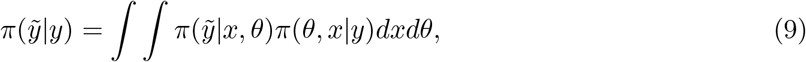

and the one step ahead out of sample predictive density is

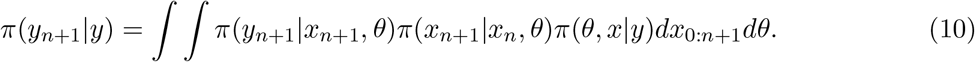

Hence, in both cases we properly account for parameter and latent process uncertainty. Although the densities in (9) and (10) will be intractable, we may generate samples via Monte Carlo, see Supplementary Information Section S2 for further details.

## 3 Results

We assume the epidemic time series (see Section 2.1) for the number of removed trees, *R*(*t*), shown in Figure 2, can be described by the compartmental SIR model with a time-varying infestation rate (see Section 2.2). The aim is to estimate the key parameters through the Bayesian inference techniques described in Section 2.3. These are the time-varying infestation rate, *β*(*t*), with corresponding stochastic noise parameter *σ* describing 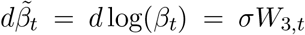, the removal rate *γ*, and the observation error *σ*_*e*_.

Section S2.4 of the Supplementary Information provides details of the assumed population sizes for each site, initial numbers of infesteds, susceptibles, and infection rate, starting parameter values for the MCMC scheme and prior specification. Regarding the latter, we take an independent prior specification for the components of *θ*, so that *π*(*θ*) = *π*(*γ*)*π*(*σ*)*π*(*σ*_*e*_). We then take lognormal *LN* (1, 1) distributions for *σ* and *σ*_*e*_, and a lognormal LN(0, 0.5^2^) distribution for *γ*. We assume that initial log infestation rate 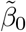 follows a Gaussian N(−8.5, 0.5^2^) distribution. These choices are motivated by the assumption of a median removal time of around 1 year (95% credible interval: (0.38, 2.66)), and a basic reproduction number at time 0 of *R*_0_ = *β*_0_*N/γ* covering a wide range of plausible values. For example, with *N* = 5 × 10^3^ the prior distributions for *γ* and 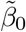 lead to a 95% credible interval for *R*_0_ of (0.25, 4.1). The initial conditions are chosen based on the increase in the removal category in the first available year, e.g., for Richmond Park there were 1414 new removals between 2013 and 2014 (new trees that had nests removed in 2014), so we assume this was approximately the number of infested locations in 2013. We investigated several choices for initial conditions and find our results robust to these variations.

### 3.1 Inference results

We ran the MCMC scheme for 10×10^3^ iterations and monitored the resulting chains for convergence. Indicative trace plots can be found in the Supplementary Information, Figure S1, and suggest that the sampler has adequately explored the parameter space. Additional chains initialised at different starting values (not shown) further confirm convergence.

From the main MCMC run we obtain the posterior within-sample means (with 50% and 95% credible intervals) for *R*(*t*), *S*(*t*) and *I*(*t*), shown in Figures 3 and 4(a-c) for Bushy and Richmond Park, respectively. The logarithmic time-dependent infestation rate, 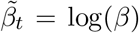, is shown in Figure 3 and 4(d). For Bushy Park, the logarithmic infestation rate is plausibly constant (given *a posteriori* variance) at 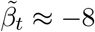, corresponding to an infestation rate of *β* = 3.4×10^−4^. Similarly, for Richmond Park, the infestation rate is plausibly constant with 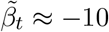, corresponding to an infestation rate of *β* = 4.5 × 10^−5^. Reassuringly, samples from the within-sample predictive for *R*(*t*) are consistent with the data used to fit the model (see panel (a) of Figures 3–4).

**Figure 3:**
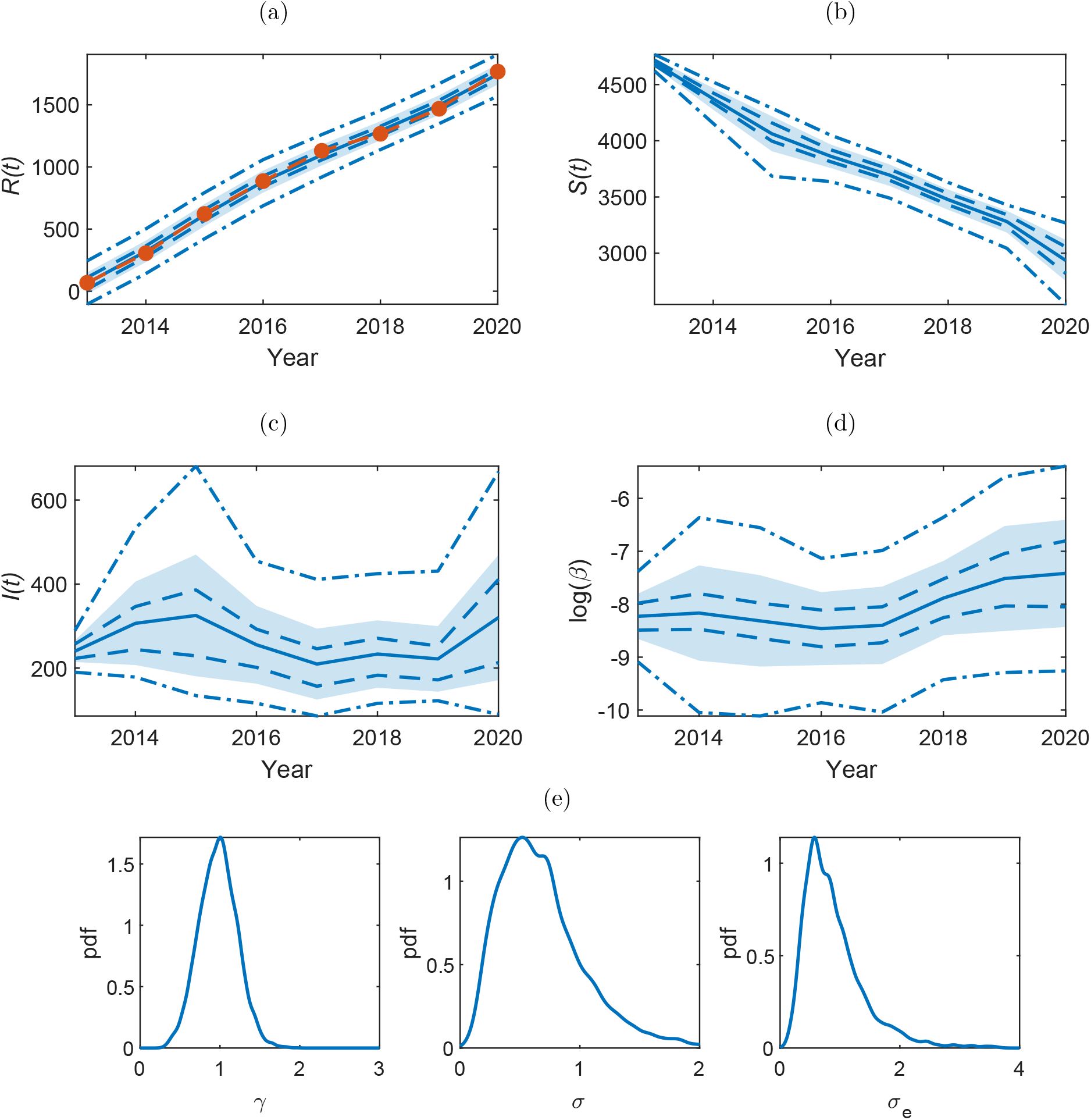
Bushy Park. The within-sample posteriors for (a) *R*(*t*), (b) *S*(*t*), (c) *I*(*t*) and (d) log(*β*_*t*_) with mean (blue solid line) ± one standard deviation (shaded region), the 50% (blue dashed), and the 95% (blue dot-dashed) credible regions. The observed time series for *R*(*t*) is overlaid in (a) (orange dashed). The corresponding (e) posterior densities for the inferred parameters *γ* (removal rate), *σ* (noise on 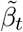) and *σ*_e_ (observation error).

**Figure 4:**
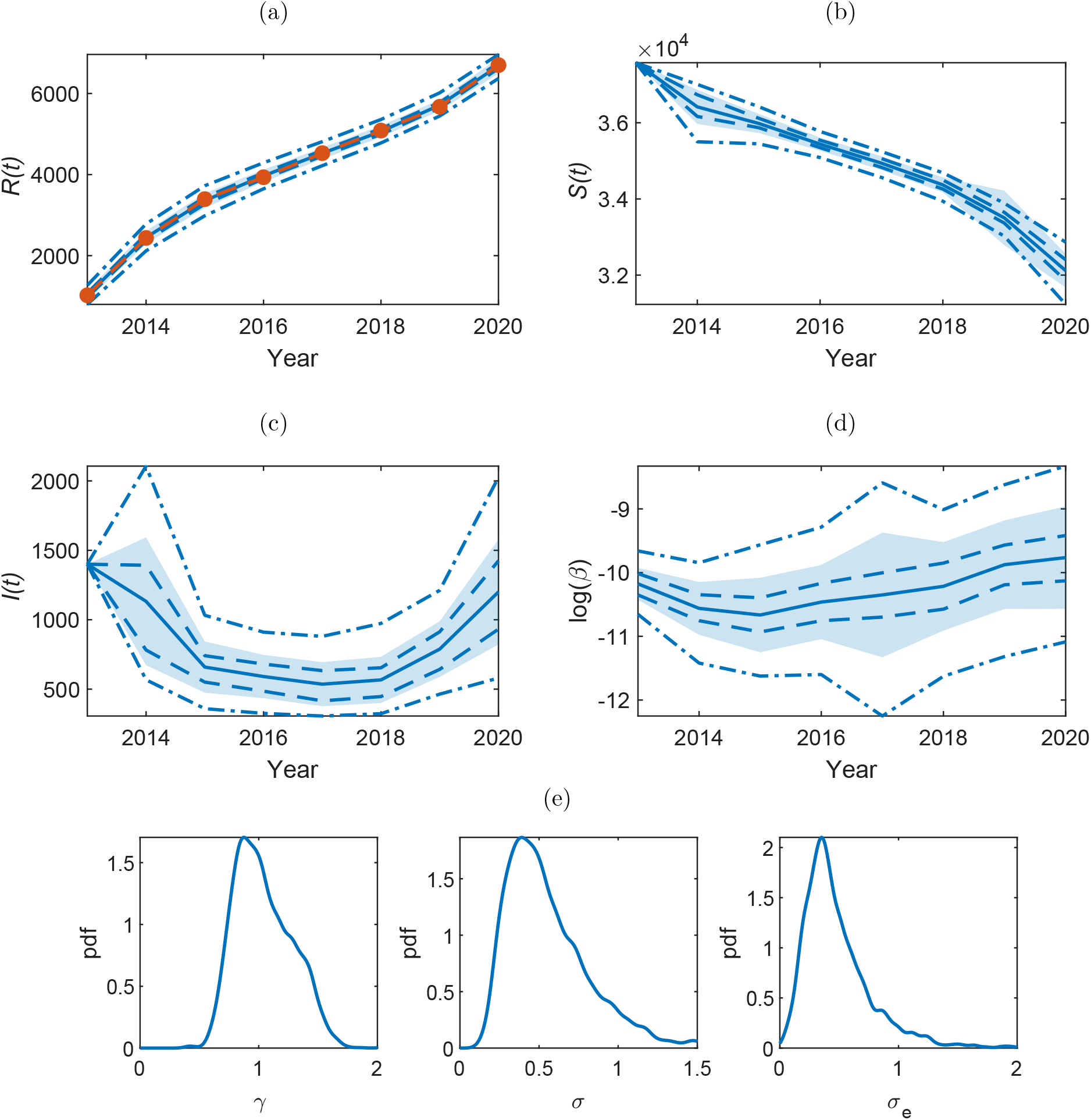
Richmond Park. The within-sample posteriors for (a) *R*(*t*), (b) *S*(*t*), (c) *I*(*t*) and (d) log(*β*_*t*_) with mean (blue solid line) ± one standard deviation (shaded region), the 50% (blue dashed), and the 95% (blue dot-dashed) credible regions. The observed time series for *R*(*t*) is overlaid in (a) (orange dashed). The corresponding (e) posterior densities for the inferred parameters *γ* (removal rate), *σ* (noise on 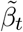) and *σ*_e_ (observation error).

The posterior density plots of the parameters *θ* = (*γ, σ, σ*_*e*_) are shown in Figures 3 and 4(e), for Bushy and Richmond park respectively. Pairwise joint posterior densities can be found in the Supplementary Information, Figure S2. The marginal posterior distribution of *γ* is centred around *γ* ≈ 1 for both Bushy and Richmond. The marginal posterior for *σ* is centred around *σ* ≈ 0.75 for Bushy and *σ* ≈ 0.5 for Richmond. The observation error *σ*_*e*_ is centred around *σ*_*e*_ ≈ 1 for Bushy and *σ*_*e*_ ≈ 0.5 for Richmond.

### 3.2 Estimation of *R*_0_

From the posterior estimations of 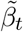 for each year and the parameter *γ*, we can estimate the basic reproduction number *R*_0_. In a deterministic system, for an epidemic to die out, *R*_0_ must be less than the threshold value of one. However, in the stochastic case it is possible for *R*_0_ to be above one but the epidemic still die out as a result of the stochastic fluctuations. Therefore it is required that *R*_0_ *<* 1 for the epidemic to shrink, upon averaging over the stochasticity. In an SIR model with a constant infection rate, *β*, the basic reproduction number is given by *R*_0_ = *βN/γ* (or simply *β/γ* if the total population number *N* has not been absorbed into the *β* constant in model formulation). Here we adapt this to use the time variant infection rate to get a reproduction number for each of the years between 2013 and 2020, *R*_0_(*t*) = *β*(*t*)*N/γ*. Box plots showing the posterior distributions of *R*_0_ for both parks are shown in Figure 5. For both parks *R*_0_ has been stable, within errors, since 2013 (corresponding to the relatively constant *β*_*t*_). However, this suggests that *R*_0_ is still above one, and therefore the epidemic will continue to propagate in these areas, and potentially beyond.

**Figure 5:**
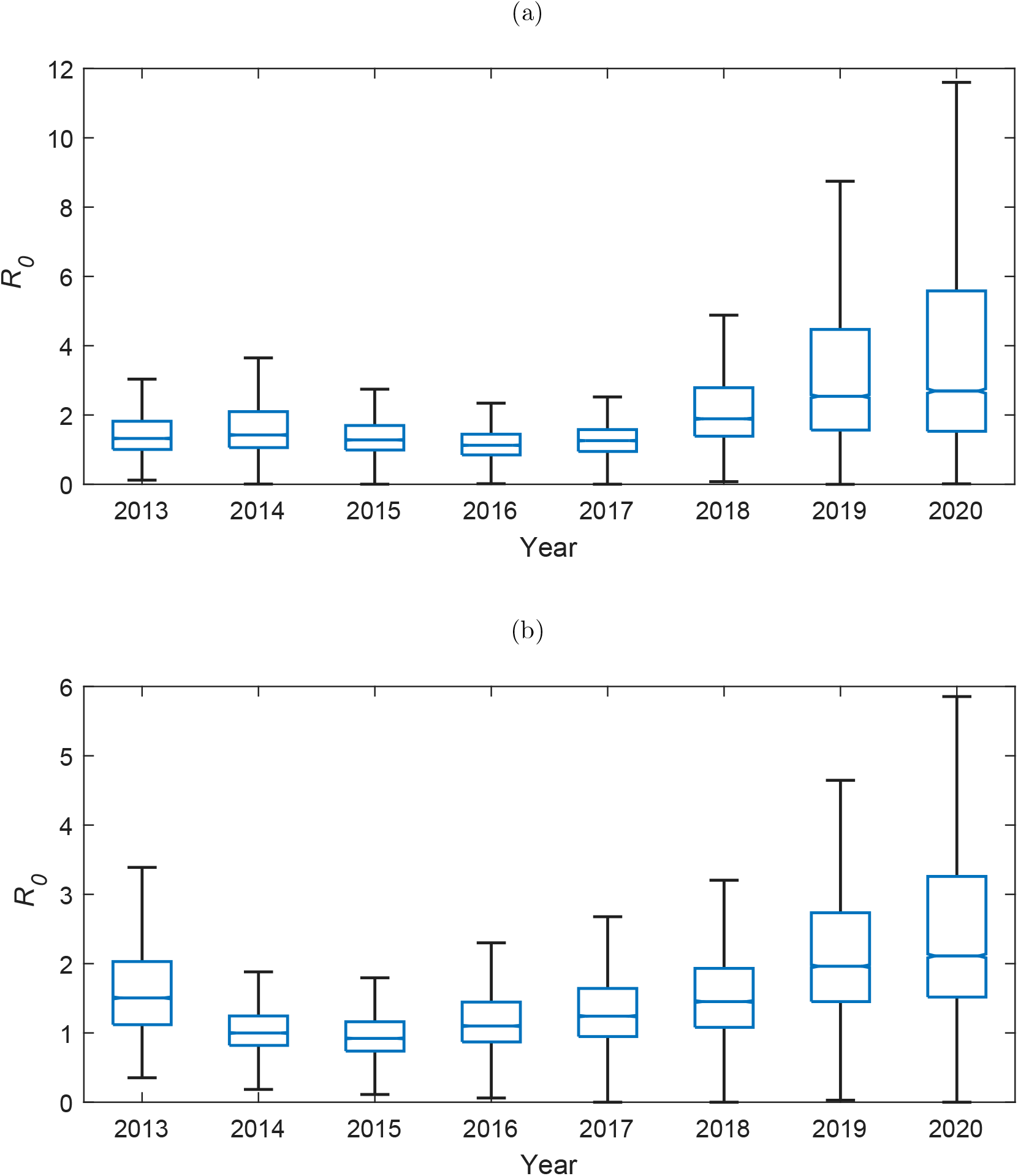
Posterior distributions of *R*_0_(*t*) = *β*_*t*_*N/γ* for (a) Bushy and (b) Richmond Park. The central line indicates the median, with the bottom and top edges of the box showing the 25th and 75th percentiles, respectively. The whiskers extend to the most extreme data points not considered outliers, which are not shown here.

### 3.3 Forward prediction

Predictions of the spread of OPM are needed to inform control strategies. To test the applicability of the SIR model with the inferred parameters and how well the model can capture future expansions in OPM, we can calculate a one year prediction. We remove the last data point, *R*(2020), and re-infer the parameters for the new shortened observed time series. We then use these parameters to run the model forwards (10 × 10^3^) simulations, matching the number of iterations in the MCMC) and obtain an estimate for *R*(2020). The median predictions with upper and lower quartiles for 1000 runs are shown in Figure 6(a) and (c) for Bushy and Richmond, respectively. In both cases, the predictive interval captures the observed data. Realisations from 100 forward runs are shown in Figure 6(b) and (d) to show that results are mostly concentrated around the observed data, with some outliers over-estimating *R*(*t*). One-step predictions for the whole time series are shown in the Supplementary Information, Figure S3.

**Figure 6:**
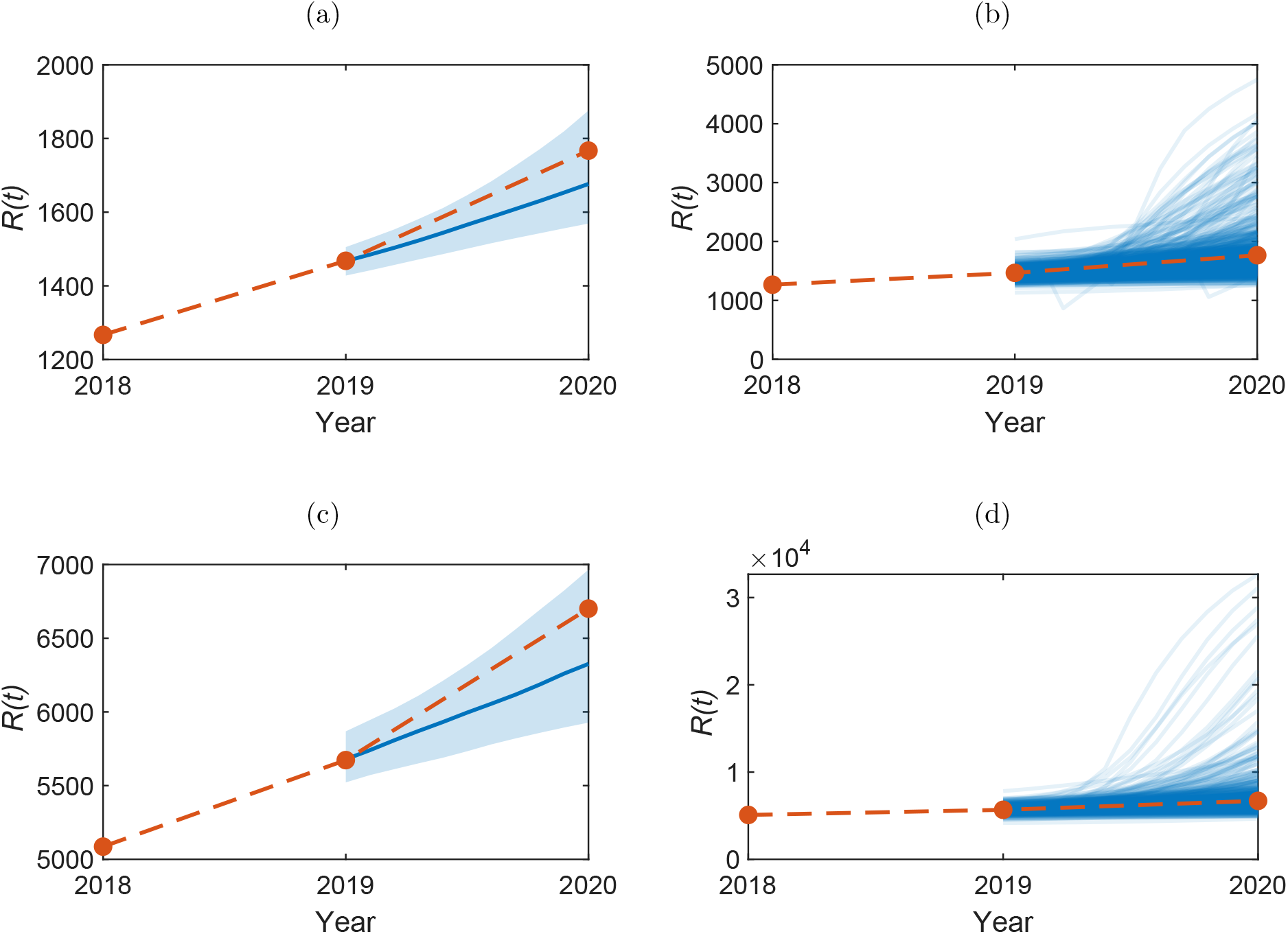
Model predictions for the number of removed nests in 2020, *R*(2020), with median (blue line) for (a-b) Bushy and (c-d) Richmond. In (a) and (c) the shaded area shows the 50% credible region. In (b) and (d) 100 simulations are shown from the forward model. The orange line shows the observed data.

Similarly we can produce predictions for the number of infested locations in 2021, *R*(2021). The median predictions with upper and lower quartiles are shown in Figure 7(a) and (b) for Bushy and Richmond, respectively. This corresponds to an average (median) of 350 new infested locations (lower-upper quartile estimate range 150–800) in Bushy Park and 1100 (700–2000) in Richmond Park.

**Figure 7:**
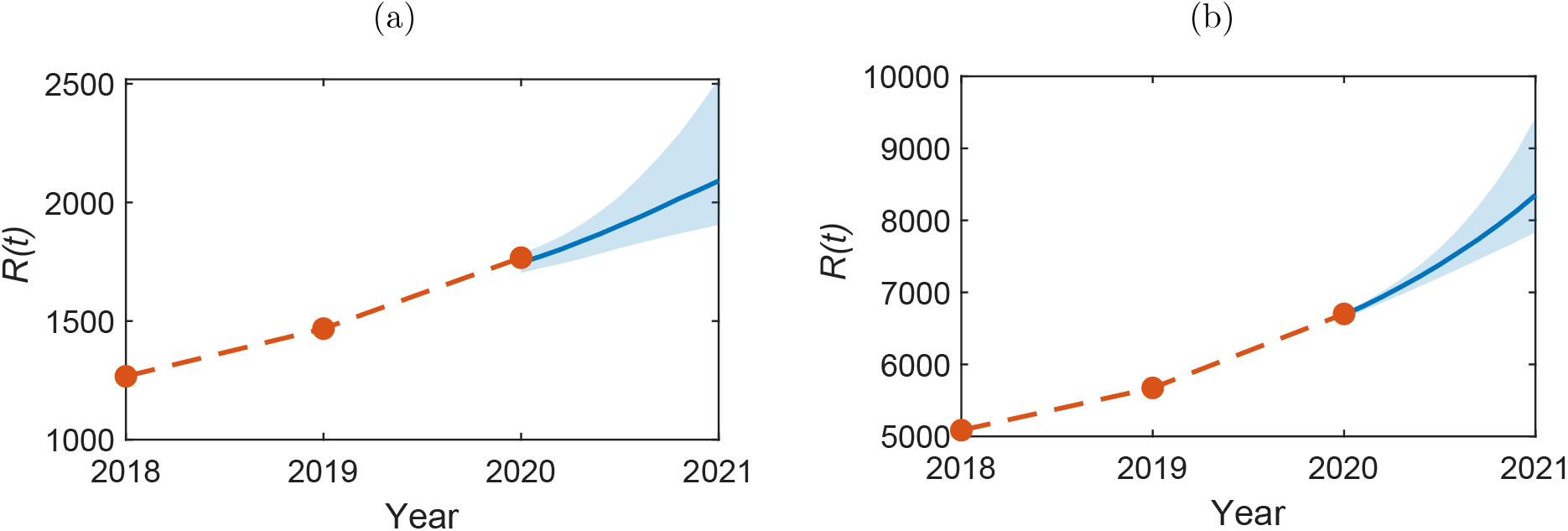
Model predictions for the number of removed nests in 2021, *R*(2021), with median (blue line) for (a) Bushy and (b) Richmond park. The shaded area shows the 50% credible region. The orange line shows the observed data up to 2020.

## Discussion

Recent modelling work has suggested that the surroundings of the current OPM infection area in the UK are highly climatically suitable and therefore at very high risk from future infestations [17]. Since the government strategy for the containment of OPM relies on targetted control at the boundary of the current infested area, it is crucial to understand and be able to predict the future spread to optimize both the cost and efficacy of these control programmes [34].

We have shown the applicability of an SIR compartmental model with a time varying infection rate to describe the OPM epidemic in the UK between the years 2013 and 2020. The statistical methodology used is a powerful tool for inferring the parameters of such models from real data and is transferable to other epidemiological and ecological datasets. Previously, similar methodology has been used to describe the spread of infectious diseases (e.g., measles [39] and Ebola [40]) and the spatial expansion of non-native plants [24], but has not yet been applied to the study of invasive insects.

Our results show, along with previous analysis [16], that the spread of OPM is continuing at a stable rate despite the current intervention methods. Correspondingly, we show that the basic reproduction number *R*_0_ has been above one since 2013. To see a reduction in the OPM population density and to protect the surrounding areas, a reduction of *R*_0_ to below one would need to be seen. Although the basic reproduction number *R*_0_ is typically used in the modelling of infectious diseases [41, 42], here it gives an analogous measure for the new infested locations caused by currently infested locations within the pest lifetime.

For simplicity and to be better described by an SIR model, we assumed that an infested location (tree) represented one removed tree regardless of how many nests were recorded as being removed from it. However, the defoliation effects and risks to human health from OPM are closely related to nest density (i.e., the numbers of nests per tree) [43]. In future work nest density could be taken into account through a nest density dependent infection rate.

A challenge of modelling OPM and other tree pests and diseases is the lack of a complete inventory oak trees in the UK, representing the susceptible population in our SIR model. This has been previously noted and highlighted as a priority for future data collection by other modelling studies [21]. It is of particular importance for future spatial models of OPM, which require an estimate of the distribution of oak trees in the areas of interest.

It is also worth noting that many areas infested with OPM have been undergoing control measures and so any inferred infestation rates represent the dynamics under these controls, rather than the inherent parameters of the uncontrolled pest spread. In Richmond and Bushy Parks, the yearly nest removal is a control measure. It would be interesting to conduct a similar analysis on a contained area that had undergone no (or different) control measures to assess the differences in the infestation rates and thus assess the efficacy of the controls. The effect of confounding factors such as the weather, difference in landscapes and presence of other pests and parasitoids, should also be investigated.

The results from this work can inform the development of future mathematical models for the spread of OPM. These models can be used to identify at-risk regions [21] and predict the distribution of OPM on a national scale. The development of these models will require further targetted data collection to obtain complete oak tree inventories, as well as data on the population numbers and locations of OPM (or indeed any other invasive insect or pathogen).

## Author contributions

LEW, NGP, AG and AWB conceptualised the project and designed the methodology. LEW analysed the data, implemented the computational models, produced the figures and led the writing of the manuscript. LEW and AG developed the inference code. JB and AH shared the OPM data and provided ecological expertise. NGP, AG and AWB acquired the funding. All authors contributed critically to the drafts and gave final approval for publication. The authors declare no conflicts of interest.

## Acknowledgements

We thank Gillian Jonusas from The Royal Parks for sharing the OPM data for Bushy and Richmond parks.

## Supplementary information

### S1 SIR Model

#### S1.1 SIR as a Markov Jump Process

Consider an SIR model in which a population of fixed size *N* is classified into compartments consisting of susceptible (*S*), infected (*I*) and removed (*R*) individuals. Let *X*_*t*_ = (*S*_*t*_, *I*_*t*_)′ denote the numbers in each state at time *t* ≥ 0 and note that *R*_*t*_ = *N* − *S*_*t*_ − *I*_*t*_ for all *t* ≥ 0. The dynamics of {*X*_*t*_, *t* ≥ 0} can be described by a Markov jump process (MJP), that is, a continuous time, discrete valued Markov process. Assuming that at most one event can occur over an infinitesimal time interval (*t, t* + Δ*t*] and that the state of the system at time *t* is *x*_*t*_ = (*s*_*t*_, *i*_*t*_)′, the MJP is characterised by probabilities of the form

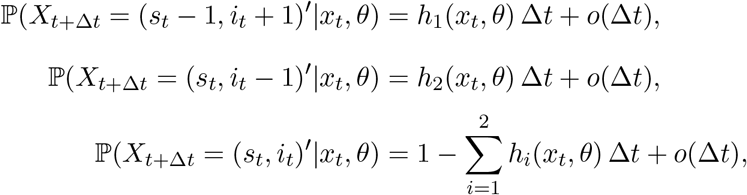

where *h*(*x*_*t*_, *θ*) = (*βs*_*t*_*i*_*t*_, *γi*_*t*_)′ is a hazard function, *θ* = (*β, γ*)′ is a parameter vector containing infection and removal rates and *o*(Δ*t*)/Δ*t* → 0 as Δ*t* → 0. The transition probability *π*(*x*_*t*_|*x*_0_, *θ*) governing the dynamics of the MJP over arbitrary time intervals of length *t* can be shown [37] to satisfy the chemical master equation (CME):

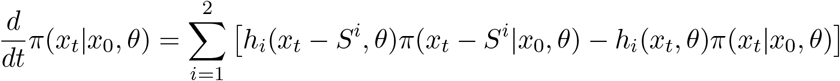

where *S*^*i*^ denotes the *i*th column of the stoichiometry matrix

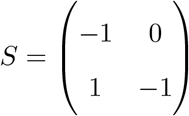

which encodes the effect of each respective transition on the components of *X*_*t*_. Although it is possible to evaluate *π*(*x*_*t*_|*x*_0_, *θ*) efficiently [44], we eschew the MJP formalism in favour of an approximation whereby *X*_*t*_ is modelled by a stochastic differential equation (SDE).

#### S1.2 SDE representation

Consider an infinitesimal time interval, (*t, t* + *dt*], over which the hazard function *h*(*x*_*t*_, *θ*) will remain constant almost surely. Let *dN*_*t*_ denote the counting process with components *dN*_1*,t*_ and *dN*_2*,t*_ containing the number of infections and removals over this interval. Hence *dN*_*i,t*_, is Poisson distributed with rate *h*_*i*_(*x*_*t*_, *θ*)*dt*. Upon noting that from *dX*_*t*_ = *SdN*_*t*_, it should be clear that

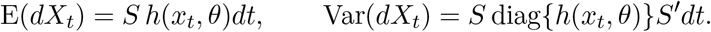

Hence, the ItÔ Stochastic differential equation (SDE) that best matches the MJP is given by

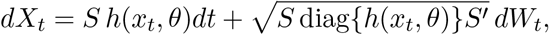

where *W*_*t*_ = (*W*_1*,t*_, *W*_2*,t*_)′ is a 2-vector of standard Brownian motion and 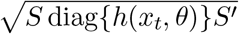 is a 2 × 2 matrix *B* such that *BB*′ = *S* diag{*h*(*x*_*t*_, *θ*)}*S*′. Explicitly, we have for the SIR model that

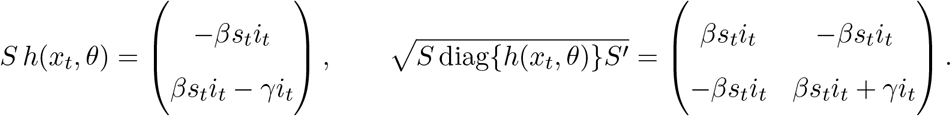

In Section 2.2 in the main text, we discuss extending the model to include a time varying infection rate.

### S2 Bayesian Inference under the LNA

We take the linear noise approximation (LNA) as described in Setion 2.2 of the main text as the inferential model. Although the observed data likelihood can be evaluated efficiently (with details below), the joint posterior is intractable. We therefore consider a Markov chain Monte Carlo scheme for generating samples from the posterior.

#### S2.1 Posterior exploration via MCMC

Our inference strategy comprises two steps:

1. Generate samples *θ*^(1)^, . . ., *θ*^(*M*)^ from the marginal parameter posterior *π*(*θ*|*y*) ∝ *π*(*θ*)*π*(*y*|*θ*).
2. Generate samples *x*^(1)^, . . ., *x*^(*M*)^ by drawing from the conditional posterior *π*(*x*|*y, θ*^(*i*)^), *i* = 1, . . ., *M*.

In step 1, we use a Metropolis-Hastings algorithm to draw from *π*(*θ*|*y*). This requires evaluation of the observed data likelihood *π*(*y*|*θ*). Note the factorisation

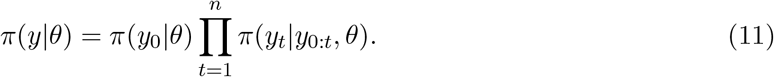

Following [32] (see also [45]), we evaluate each constituent term in (11) via a forward filter. The forward filter requires the observation equation (main text (7)) to have a linear Gaussian structure, which is not the case as written, due to the variance being a function of the latent state process. Therefore, we run step *t* of the forward filter with *P*′*x*_*t*+1_*P* replaced by *P*′*η*_*t*+1_*P*. That is, the unknown *x*_*t*+1_ is replaced by the LNA predictive mean *η*_*t*+1_.

Since the parameters *θ* remain fixed throughout the calculation of *π*(*y*|*θ*), we drop them from the notation where possible. Define *y*_0:*t*_ = (*y*_0_, . . ., *y*_*t*_)′. Now suppose that *X*_0_ ~ N(a_0_, C_0_) *a priori*. Algorithm 1 (Supplementary Information Section S2.3) gives the forward filter. This can then be used inside Algorithm 2 (Section S2.3), which uses a random walk Metropolis algorithm to generate (dependent) draws from the marginal parameter posterior *π*(*θ*|*y*). The proposal mechanism requires an innovation variance Ω which can be chosen to maximise mixing efficiency, as measured by say effective sample size per second. We take 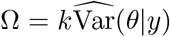 where the posterior variance is estimated from a short pilot run and *k* is chosen subsequently to give an acceptance rate of around 20–30% [46].

Given samples *θ*^(1)^, . . ., *θ*^(*M*)^ from *π*(*θ*|*y*), we generate samples of the latent process *x*^(*i*)^ ~ *π*(*x*|*y, θ*^(*i*)^), *i* = 1, . . ., *M* by noting that these samples can be efficiently generated using a backward sampling algorithm. This requires the covariance between *X*_*t*_ and *X*_*t*+1_ (since the former is drawn conditionally on a realisation of the latter) which depends on the 3 × 3 fundamental matrix *G*_*t*_. This satisfies and ODE system

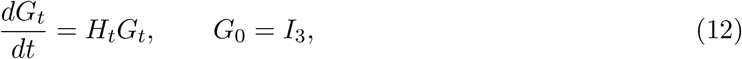

where *I*_3_ is the 3 × 3 identity matrix. The ODE in (12) can be time-stepped with the ODEs in (4) and (6) (main text) as part of the forward filter. Algorithm 3 (Section S2.3) then gives the backward sampler.

#### S2.2 Model checking

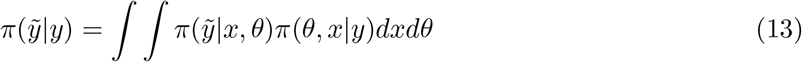

and the one step ahead out of sample predictive density is

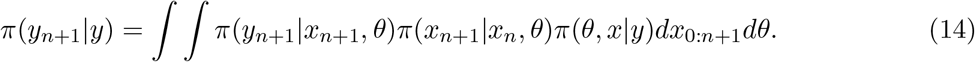

Hence, in both cases we properly account for parameter and latent process uncertainty. Although the densities in (13) and (14) will be intractable, we may generate samples via Monte Carlo. Recall that the inference algorithm described in Section S2.1 gives draws {(*θ*^(*i*)^, *x*^(*i*)^), *i* = 1, . . ., *M*}. We then generate 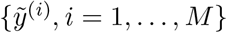 via (7), by drawing 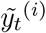 from a 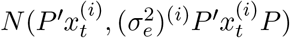 distribution, independently for *t* = 0, . . ., *n* and *i* = 1, . . ., *M*. Similarly, we generate samples from *π*(*y*_*n*+1_|*y*) by first drawing 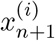 from 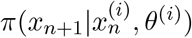, followed by 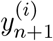 from 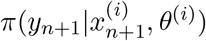.

#### S2.3 Algorithms

**Algorithm 1.**
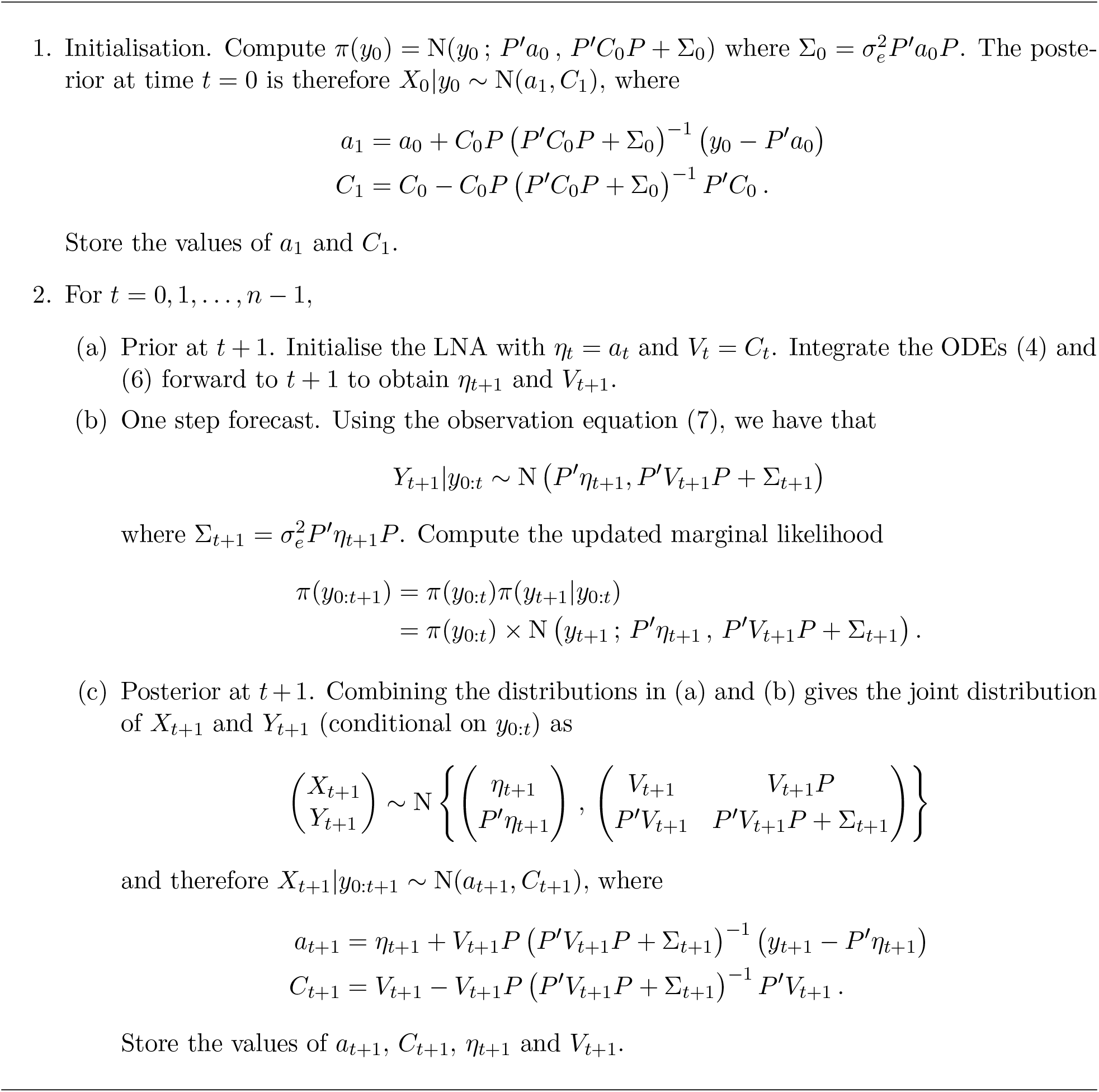
LNA forward filter

**Algorithm 2.**
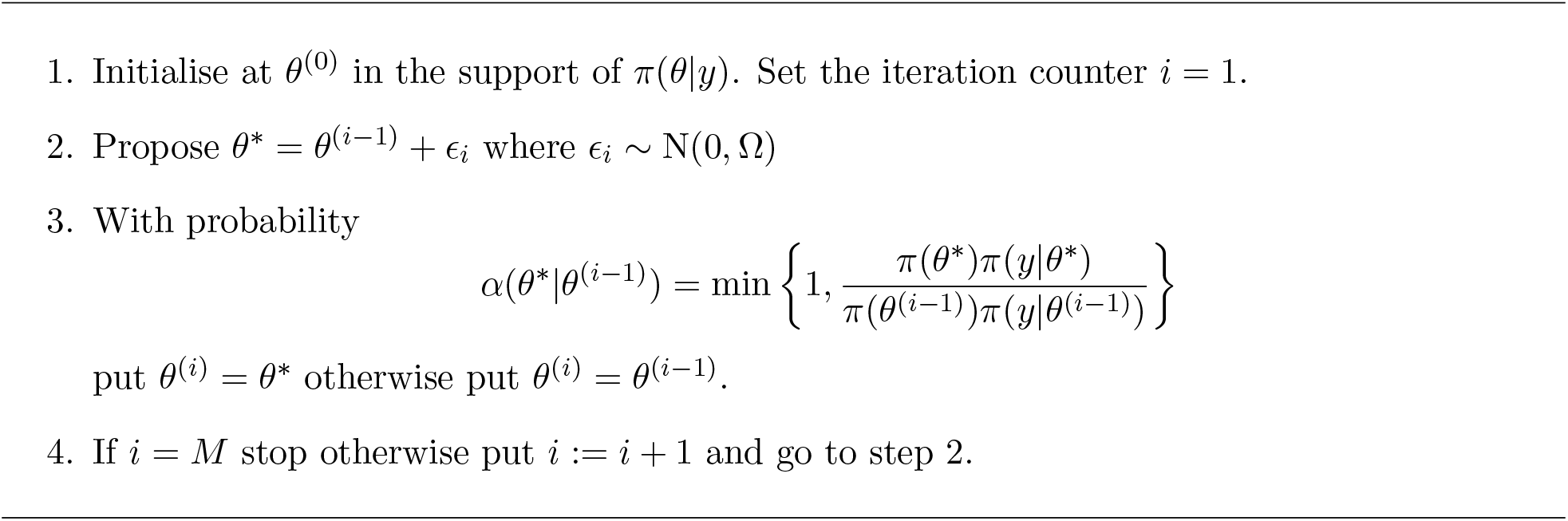
Random walk Metropolis algorithm

**Algorithm 3.**
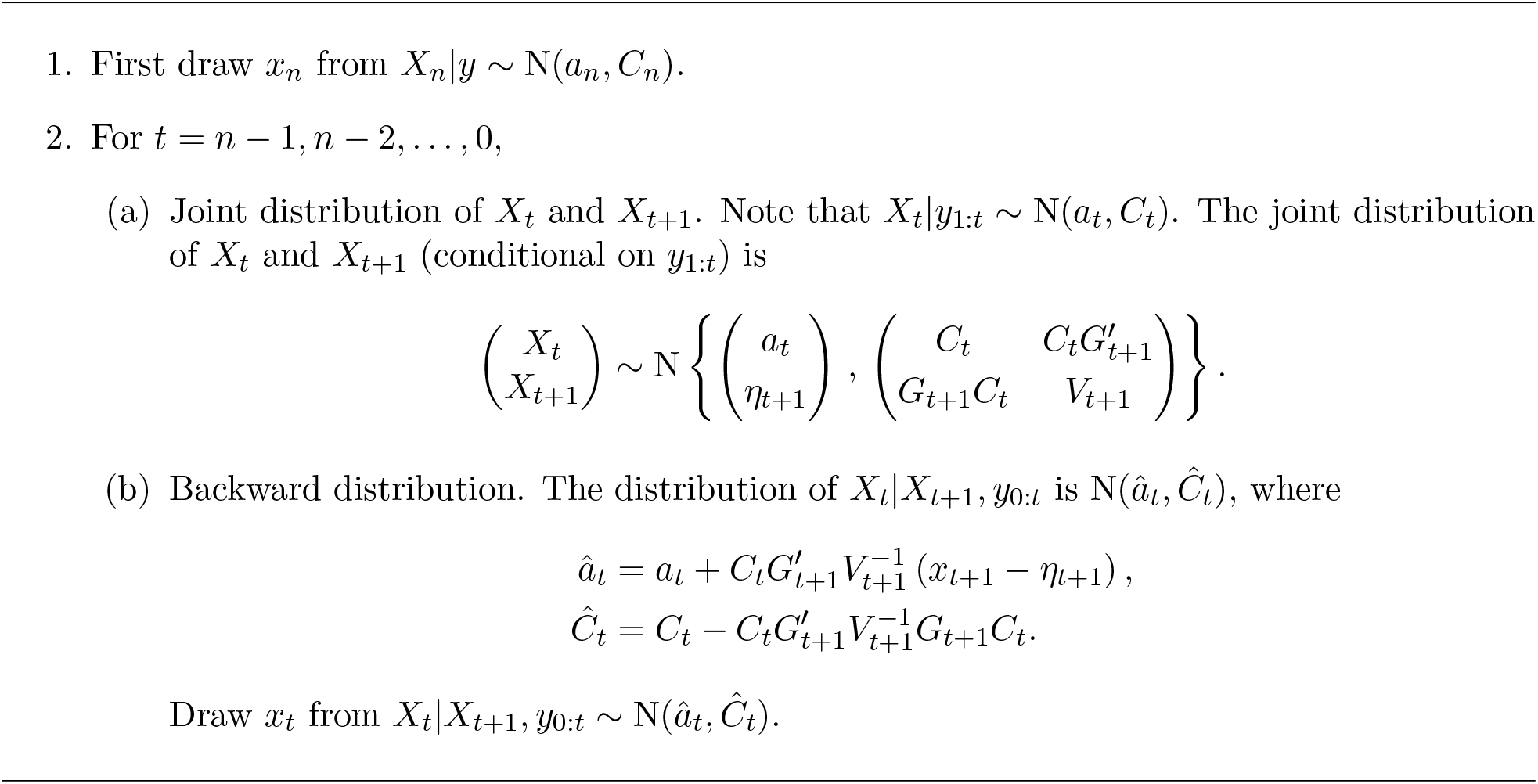
LNA backward sampler

#### S2.4 Parameters for inference

**Table S1:** The parameters used in the inference schemes described in Sections 2.3 and S2.

|  |  | Bushy | Richmond |
| --- | --- | --- | --- |
| Total population | $N$ | 5000 | 40000 |
| Initial infected | $I_0$ | 240 | 1400 |
| Initial infection rate | $\tilde{\beta}_0$ | -8.5 | -10 |
| Initial model parameters | $\theta_0 = (\gamma_0, \sigma_0, \sigma_e)'$ | $(1, 0.5, 1)'$ | $(2, 0.5, 1)'$ |
| Observation matrix | $P$ | $(1, 1, 0)$ | |
| Prior distributions | $\pi(\theta)$ | $\log \gamma \sim N(0, 0.5^2)$ | |
| | | $\log \sigma \sim N(1, 1)$ | |
| | | $\log \sigma_e \sim N(1, 1)$ | |
| Tuning parameter | $\Sigma$ | $\begin{pmatrix} 0.1 & 0 & 0 \\ 0 & 0.1 & 0 \\ 0 & 0 & 1 \end{pmatrix}$ | |
| Initial conditions mean | $a_0$ | $(N - I_0 - R_0, I_0, \tilde{\beta}_0)$ | |
| Initial conditions variance | $C_0$ | $\begin{pmatrix} 0 & 0 & 0 \\ 0 & 0 & 0 \\ 0 & 0 & 0.5 \end{pmatrix}$ | |

### S3 Additional results figures

The following section contains additional results figures referred to in the main text. We give the trace (Figure S1) and 2D contour plots (Figure S2) for the inferred model parameters. The one-step model predictions for all years are given in Figure S3

**Figure S1:**
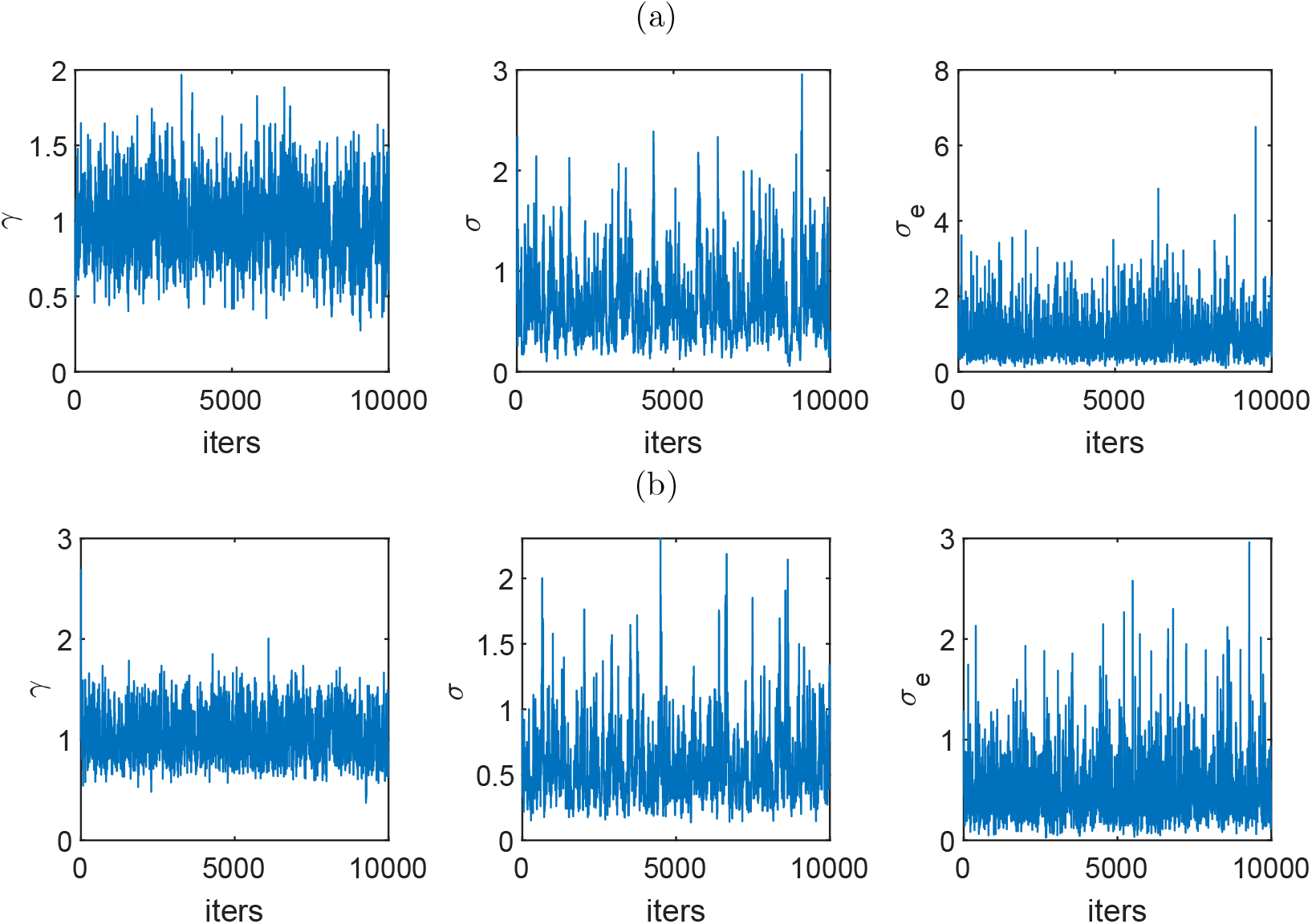
Trace plots for (a) Bushy and (b) Richmond Park.

**Figure S2:**
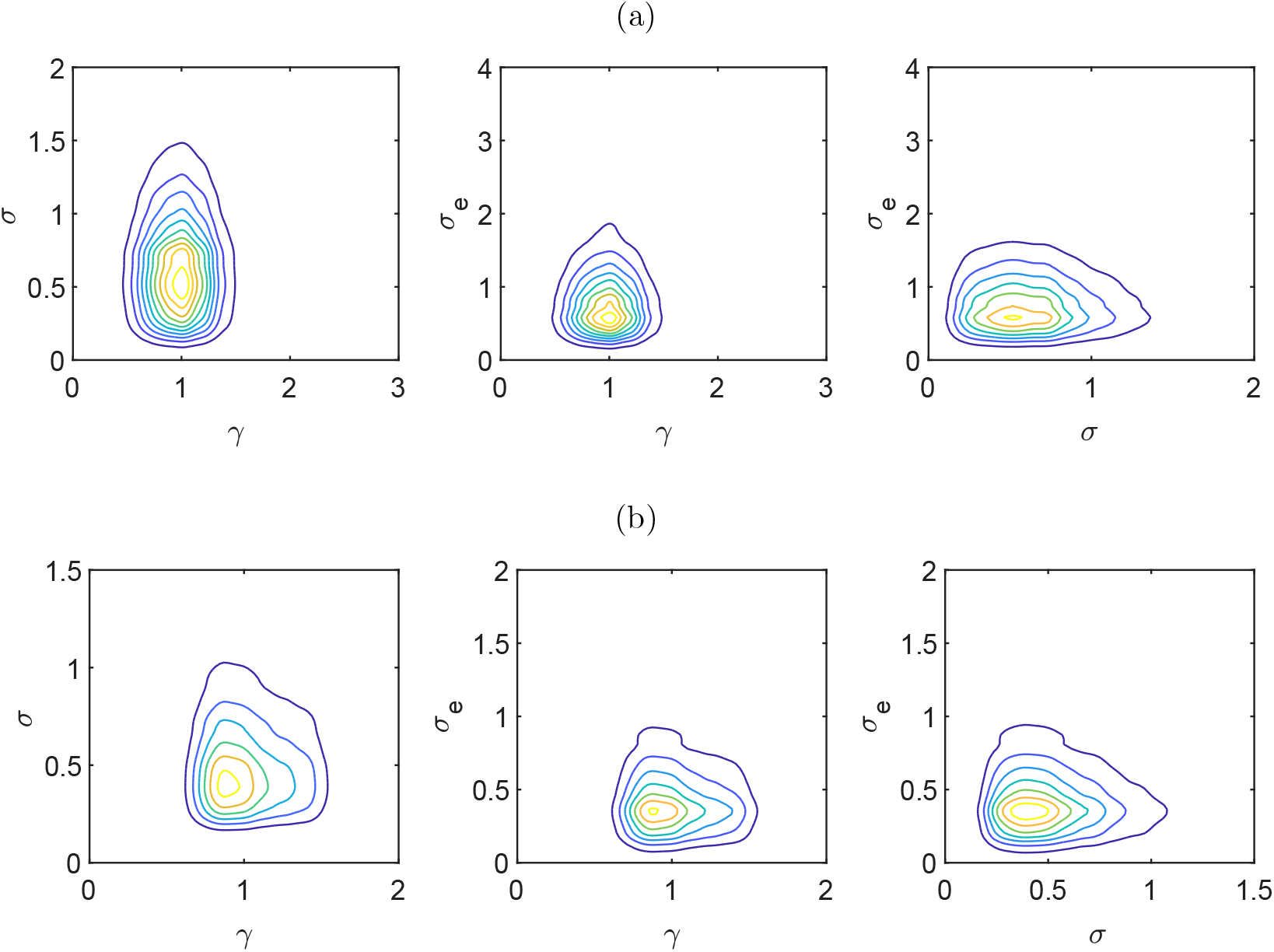
Pairwise joint posterior densities for the parameters *γ*, *σ* and *σ*_*e*_ for (a) Bushy and (b) Richmond Park.

**Figure S3:**
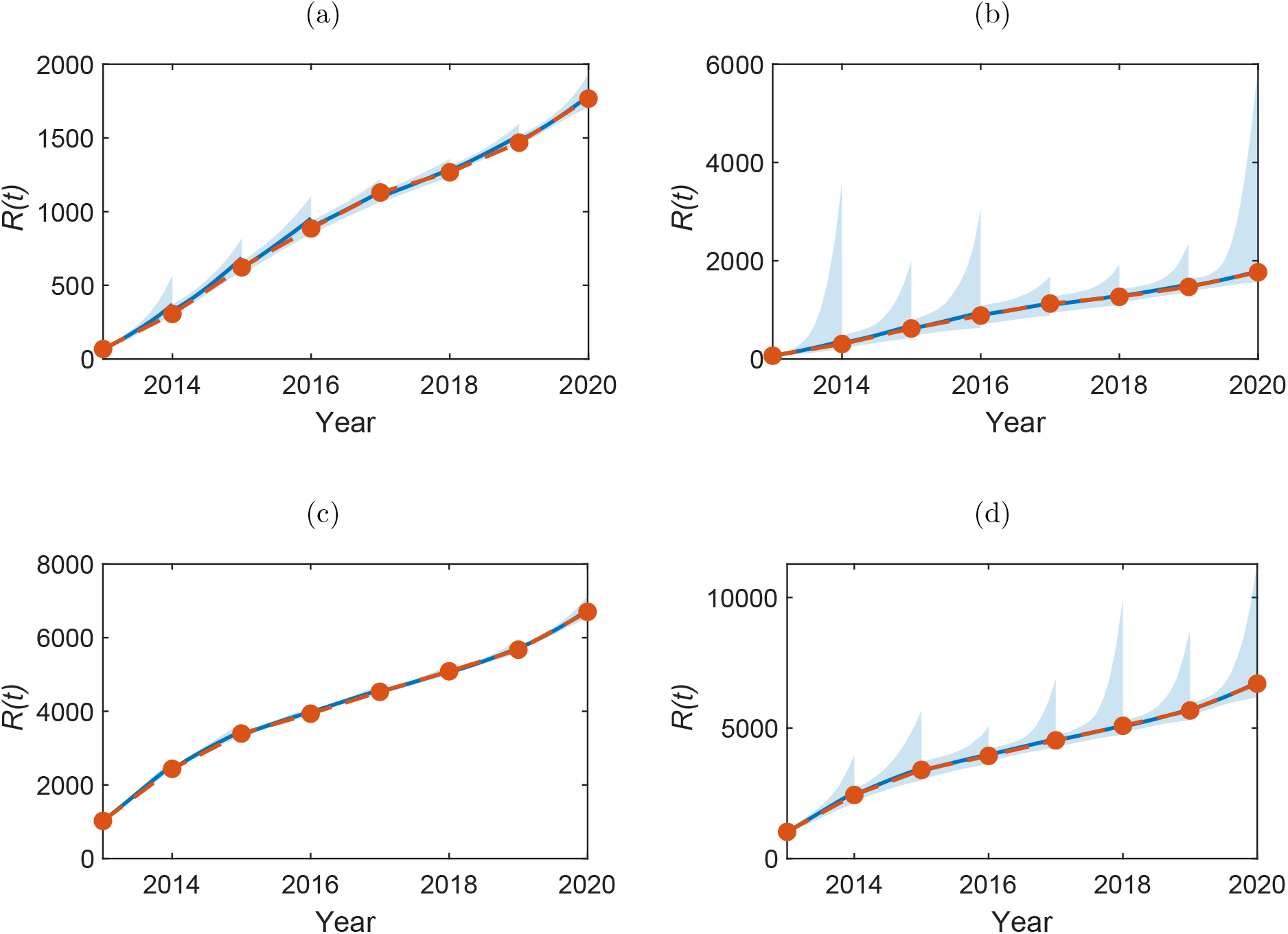
One-step model predictions for the number of removed nests, *R*(*t*) for the years 2014– 2020, with median (blue line) for (a-b) Bushy and (c-d) Richmond. In (a) and (c) the shaded area shows 50% credible region and in (b) and (d) the 95% credible region. The orange line shows the observed data.

